# Glycolytic reliance promotes anabolism in photoreceptors

**DOI:** 10.1101/101964

**Authors:** Yashodhan Chinchore, Tedi Begaj, David Wu, Eugene Drokhlyansky, Constance L. Cepko

## Abstract

Sensory neurons capture information from the environment and convert it into signals that can greatly impact the survival of an organism. These systems are thus under heavy selective pressure, including for the most efficient use of energy to support their sensitivity and efficiency^1^. In this regard, the vertebrate photoreceptor cells face a dual challenge. They not only need to preserve their membrane excitability via ion pumps by ATP hydrolysis^2^ but also maintain a highly membrane rich organelle, the outer segment, which is the primary site of phototransduction, creating a considerable biosynthetic demand. How photoreceptors manage carbon allocation to balance their catabolic and anabolic demands is poorly understood. One metabolic feature of the retina is its ability to convert the majority of its glucose into lactate^3,4^ even in the presence of oxygen. This phenomenon, aerobic glycolysis, is found in cancer and proliferating cells, and is thought to promote biomass buildup to sustain proliferation^5,6^. The purpose of aerobic glycolysis in the retina, its relevance to photoreceptor physiology, and its regulation, are not understood. Here, we show that rod photoreceptors rely on glycolysis for their outer segment (OS) biogenesis. Genetic perturbations targeting allostery or key regulatory nodes in the glycolytic pathway impacted the OS size. Fibroblast growth factor (FGF) signaling was found to regulate glycolysis, with antagonism of this pathway resulting in anabolic deficits. These data demonstrate the cell autonomous role of the glycolytic pathway in OS maintenance and provide evidence that aerobic glycolysis is part of a metabolic program that supports the biosynthetic needs of a normal neuronal cell type.

---

Photoreceptor cells maintain elaborate membranous organelles (rhabdomeres in invertebrates and OS in vertebrates) that maximize light capture. The maintenance of such structures requires considerable metabolic resources. Invertebrate photoreceptors exhibit light-dependent endocytosis of their photosensitive membranes^7^ enabling the recycling of these resources. In contrast, vertebrate photoreceptors shed a fraction of their OS daily, to be phagocytosed by the juxtaposed retinal pigmented epithelium (RPE)^8,9^ (Extended Data Fig. 1). To sustain a constant volume of the OS, a cell must channel metabolites towards biosynthesis, against the backdrop of very high ATP consumption, which is required to maintain membrane potential. Photoreceptors thus must balance the use of their intracellular carbon pool between oxidative catabolism, to generate the required ATP, and anabolism, to continually renew the OS.

**Figure 1.**
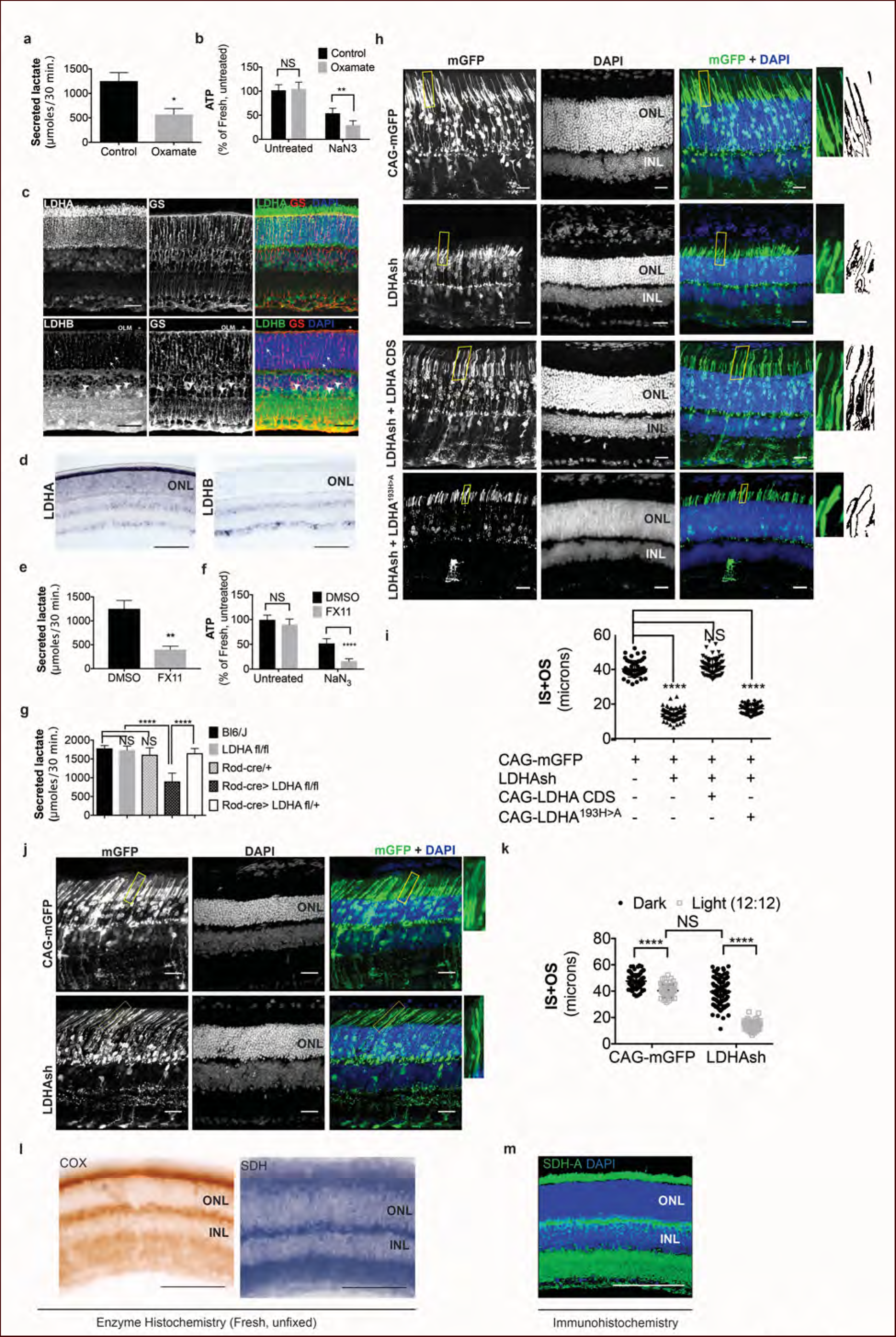
*LDHA*-dependent aerobic glycolysis and outer segment maintenance in photoreceptors **a**. Freshly explanted retinas were treated with the Ldh inhibitor, sodium oxamate, for 8 h, transferred to Krebs’ Ringers for 30’, and lactate was measured in the supernatant. Control (n=4), Oxamate (n=5). **b**. Freshly explanted retinas were treated with oxamate or NaCl (control) in medium for 8 h, followed by treatment with NaN3 or NaCl (untreated group) in Krebs’ Ringers medium for 30 minutes. ATP per retina was then measured. *n*=7, Control untreated; *n*=8, Oxamate untreated *n*=8, Control NaN_3_; *n*=8, Oxamate NaN_3_. **c**, Expression of *LDHA* and *LDHB* as determined by IHC. Glutamine synthetase (GS), a Mueller glia-specific marker, colocalized with LDHB in the cell bodies (arrowheads), processes ensheathing the photoreceptors (arrows) and the outer limiting membrane (OLM, ^*^). Scale bar, 50 μm. **d**, ISH for *LDHA* and *LDHB*. *LDHA* RNA displayed photoreceptor-enriched expression while *LDHB* RNA was not observed in photoreceptors. Scale bar, 100 μm. **e,f** Freshly explanted retinas were treated with FX11 or DMSO for 8 hours and transferred to Krebs’ Ringers for 30 minutes and secreted lactate was measured (e), or they were transferred to Krebs’ Ringers buffer with NaN3 or NaCl (untreated group) for 30 minutes for ATP quantitation (f). ATP per retina was measured at the end of the assay. *n*=8, DMSO untreated; *n*=8, FX11 untreated; *n*=9, DMSO NaN_3_; *n*=7, FX11 NaN_3_. **g**, Freshly explanted retinae were transferred to Krebs’ Ringers for 30 minutes and secreted lactate was measured. *n*=8, Bl6/J; *n*=8, *LDHA*^*fl/fl*^; *n*=8, Rod-cre; *n*=16, Rod-cre> *LDHA*^*fl/fl*^*; n*= 8, Rod-cre> *LDHA*^*fl/fl*^. **h**, Photoreceptor outer segment phenotype 42-45 days following *in vivo* electroporation of a knock-down construct (shRNA) for LDHA. CAG-mGFP was used for coelectroporation. Plasmid combinations listed on the left. Magnification of areas outlined in yellow is displayed on right with threshold-adjusted rendering to highlight inner and outer segments. Scale bar, 25 μm. **i**, Quantification of inner+outer segment (IS+OS) lengths. n= 53-74 photoreceptors, 4-5 retinae. **j**, Photoreceptor outer segment phenotype of dark-reared animals. Electroporated pups were transferred to dark on the day of eye opening (P11) and reared with their mothers for 3 weeks. **k**, Quantification of inner+outer segment lengths of (j). n= 53-83 photoreceptors, 4-5 retinae. **l**, Colored end products of redox reactions catalyzed by COX and SDH enzymes in retinal tissue. Scale bar, 200 μm. **m**, IHC for SDH-A subunit in adult retina. Scale bar, 200 μm. ONL, outer nuclear layer. INL, inner nuclear layer. Data, Mean±SD. Statistics, unpaired, two-tailed *t*-test with Kolmogorov-Smirnov correction for panels a, e; two-way ANOVA with Tukey’s correction for panels **b, f** and **k**; one-way ANOVA with Tukey’s multiple comparison test for panels **g**, **i**.

The mammalian retina depends upon glucose and glycolysis for survival and function^10,11^. The majority (~80%) of glucose is converted to lactate via glycolysis^3,12^. The adult retina can produce lactate aerobically (aerobic glycolysis/Warburg effect) with an ~50% increase during anaerobic conditions (Pasteur effect)^4^. The cell types that carry out aerobic glycolysis in the normal adult retina have not been determined. The photoreceptors have been assumed to rely on aerobic glycolysis. This assumption is based on the adverse effects on photoreceptor function after *en masse* inhibition of whole retinal glycolysis by pharmacological treatments e.g. with iodoacetate^12^. The Warburg effect exemplifies an elaborate set of metabolic strategies adopted by a cell to preferentially promote glycolysis^5,6^. One drawback of inhibiting glycolytic enzyme activity to ascertain its role in the functions of the retina is that such a manipulation does not differentiate between aerobic glycolysis and housekeeping glycolysis- a pathway critical for most cell types.

Studies conducted on retinal tissue *in vitro* indicate that isolated mammalian photoreceptors can consume lactate, which can be produced by glycolysis in retinal Mueller glia^13^. Thus the decreased photoreceptor function after whole retinal glycolytic enzyme inhibition could be a non cell-autonomous effect on Muller glia. Though many features of the “lactate shuttle” and its *in vivo* relevance have recently been questioned^14^, it is important to devise an experimental strategy that would be able to discern the cell-autonomous versus non-autonomous requirement of glycolysis for the photoreceptors.

We first examined lactate production from the retina and assayed the metabolic consequences of inhibiting aerobic glycolysis. Lactate is produced by reduction of pyruvate, a reaction catalyzed by lactate dehydrogenase (Ldh) (Extended Data Fig. 2a). Freshly isolated retinae were cultured in the presence or absence of sodium oxamate- an Ldh inhibitor. These were subsequently transferred to buffered Krebs’-Ringer’s medium that has glucose as the sole source of carbon (see Methods), and lactate secretion was quantified (Fig. 1a). The extracellular secreted lactate was measured because it represents the pyruvate-derived carbons that are diverted away from other intracellular metabolic processes or the mitochondria. In addition, the ATP levels were monitored and, surprisingly, the steady state levels of ATP in oxamate-treated retinae did not differ from the control retinae (Fig. 1b). This could be due to a relatively minor glycolytic contribution to the total ATP pool, a compensatory metabolic realignment towards mitochondria-dependent ATP production or adenylate kinase-dependent ATP synthesis. To differentiate among these possibilities, explants were cultured in oxamate or control conditions followed by a short treatment with NaN_3_ to inhibit mitochondrial ATP synthesis (Fig. 1b). Control retinae displayed ~50% reduction in ATP levels after incubation in NaN_3_. Interestingly, oxamate-treated retinae displayed a further 20% decrease in ATP after exposure to NaN_3_. Thus, inhibiting lactate synthesis resulted in a greater fraction of the ATP pool that was sensitive to mitochondrial function.

**Figure 2.**
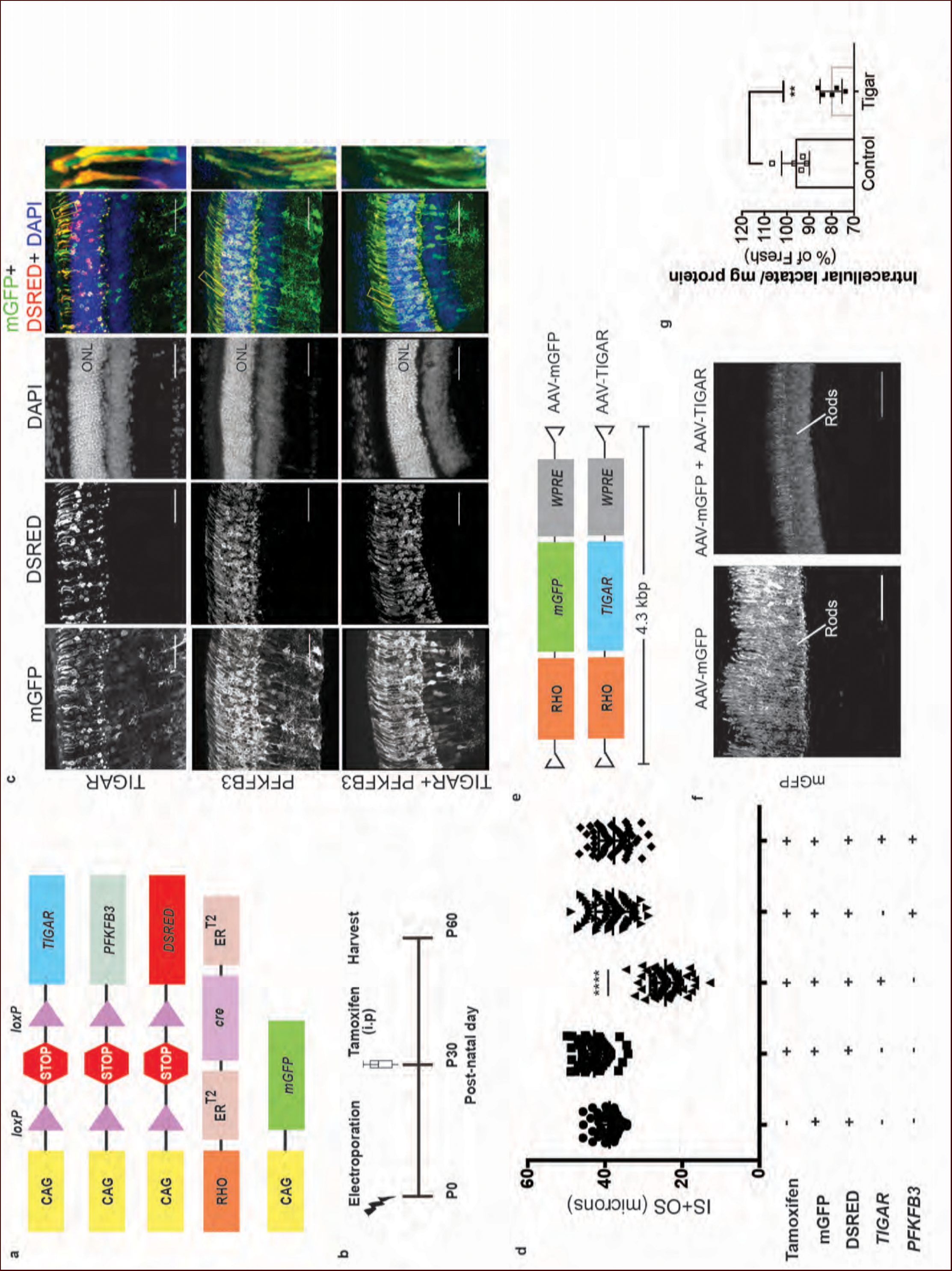
Targeting allostery reveals glycolytic reliance for outer segment maintenance. **a**, Constructs for spatio-temporal control of expression of *TIGAR* and *PFKFB3*. DsRed used as the *cre* reporter, mGFP as a coelectroporation marker. **b**, Scheme for electroporation and tamoxifen induction. i.p, Intraperitoneal. **c,d**, IS+OS length were measured following introduction of *TIGAR* (n=72 cells), *PFKB3* (n=72 cells), and *TIGAR* and *PFKFB3* constructs (n=78), shown in (a). Controls were -tamoxifen (n=62) and +tamoxifen (n= 74). Scale bar, 50 μm. Data are Mean±SD. One-way ANOVA with Tukey’s correction. Outlined areas magnified to show IS and OS morphology. **e**, AAV genomes for expression of mGFP (AAV-mGFP) or *TIGAR* (AAV-TIGAR). **f**, Cross sections of AAV-mGFP or AAV-TIGAR infected retinae harvested at P28 imaged for mGFP expression. **g**, Intracellular lactate normalized for total protein was quantified for retinae infected with AAV-mGFP (Control) or AAV-mGFP + AAV-TIGAR. Data represented as percentages relative to age-matched, freshly isolated retinae. Data, Mean±SD. Unpaired, two-tailed *t*-test with Kolmogorov-Smirnov correction for non-Gaussian distribution. ONL, outer nuclear layer.

Next, we wanted to ascertain if photoreceptors produce or consume lactate. As a first step, the expression of the Ldh subtypes was examined. Ldh is a tetrameric enzyme composed of Ldha and Ldhb subunits encoded by the *LDHA* and *LDHB* genes respectively. The subunits can assemble in five different combinations with differing kinetic properties^15,16^ (Extended Data Fig. 2b). A tetramer of all Ldha subunits has high affinity for pyruvate and a higher *V_max_* for pyruvate reduction to lactate. In addition, many glycolytic cancers have elevated *LDHA* expression^17,18^. On the contrary, an all-Ldhb tetramer is maximally active at low pyruvate concentrations, is strongly inhibited by pyruvate and is expressed in tissues using lactate for oxidative metabolism or gluconeogenesis^15^. We examined the expression of Ldha and Ldhb subunits in the retina by immunohistochemistry (IHC) (Fig. 1c). Photoreceptors showed strong expression of Ldha, particularly with respect to the other retinal cell types. Ldhb was abundantly expressed in the cells of the inner nuclear layer (INL), which includes interneurons and Mueller glia. Expression analysis of transcripts of *LDHA* and *LDHB* genes by *in situ* hybridization (ISH) (Fig. 1d) confirmed that *LDHA* is enriched in the photoreceptors, whereas *LDHB* is excluded. This was also confirmed by qRT-PCR analysis of *LDHA* and *LDHB* transcripts in isolated rod photoreceptor cDNA (Extended Data Table 1). We conclude that photoreceptors have predominantly Ldha-type subunits.

We also assessed the ability of the retina to secrete lactate after treatment with the Ldha-specific inhibitor, FX-11^19^. FX-11 significantly reduced lactate secretion (Fig. 1e). Similar to oxamate, FX-11 also resulted in an increased percentage of ATP that was sensitive to azide inhibition (Fig. 1f). To investigate if photoreceptors produce lactate in an *LDHA*-dependent manner, mice with a conditional allele of *LDHA*^20^, (*LDHA*^*fl/fl*^), were used. The specificity and efficiency of - Cre recombinase under control of the rhodopsin regulatory elements (Rod-cre)^21^ were first tested, which showed that only rod photoreceptors had a history of *cre* expression (Extended Data Fig. 2c, d). The recombination efficiency varied between ~50%-90% of photoreceptors among different retinae. The Rod-cre/*LDHA*^*fl/fl*^ retinae were examined for Ldha protein, which showed a significant reduction (Extended Data Fig. 2e). A compensatory expression of *LDHB* in photoreceptors was not detected (Extended Data Fig. 2f). Lactate production in these retinae was examined and was found to be significantly reduced (Fig. 1g). Thus photoreceptors produce lactate in an *LDHA-*dependent manner.

In order to assess if reduction in *LDHA* created a cellular phenotype in photoreceptors, and if so, whether it was required autonomously, electroporation of a short hairpin RNA (shRNA) specifically targeting the 3’ untranslated region (UTR) of the *LDHA* transcript was used (Extended Data Fig. 2g). This strategy was taken, vs. examination of the rods in the Rod-cre/*LDHA*^*fl/l,*^ retinae, due to the concern that a reduction in lactate by rods might affect closely associated cell types, such as Mueller glia and/or RPE cells, creating non-autonomous effects on rods. A plasmid encoding this shRNA was delivered to the retina *in vivo* by electroporation. Electroporation occurs in patches comprising 15-30% of the retina, and in a given patch, only ~20% cells are electroporated (Sui Wang and C. Cepko, unpublished). Thus, plasmid transfection via electroporation allowed us to determine if *LDHA* has a cell-autonomous role in photoreceptors. The electroporated photoreceptors had markedly reduced OS length when compared to control (Fig. 1h,i). Genetic complementation by coelectroporation of a sh-resistant *LDHA* cDNA that lacks the 3’UTR demonstrated that the defect was attributable to *LDHA* loss-of-function (Fig. 1h,i). To determine if the catalytic activity of Ldha was required for rescue, an allele of *LDHA* with a point mutation in the catalytic center (LDHA^H193>A^) was introduced. It failed to rescue the shRNA phenotype. Finally, expression of *LDHB* was not sufficient to compensate for *LDHA* loss-of-function (Extended Data Fig. 2h).

The cyclical process of OS shedding and renewal is regulated by light^9^. Since Ldha function is necessary to maintain OS length, we assessed the effect of *LDHA* knockdown in dark-reared mice and compared with mice raised in normal room light. Electroporated mice were raised with their mothers in normal room light until eyes were open (P11), and then shifted to the dark for 3 weeks. In mice with no *LDHA* knockdown, there was ~25% increase in OS length after dark rearing compared to light/dark rearing (Fig. 1j, k), presumably as a result of less OS shedding. Interestingly, in mice with the *LDHA* knockdown, dark rearing resulted in loss of the LDHA knockdown phenotype (Fig. 1j, k). The average length of *LDHA* knockdown OS was similar to that of light-treated control animals. These data indicate that reducing the need for OS biogenesis, as occurs in the dark, led to a reduced reliance on *LDHA* function.

Cells with immature or dysfunctional mitochondria become reliant on glycolysis by increasing *LDHA* expression at the expense of *LDHB*^22,23,24^. Though photoreceptors have abundant mitochondria, a reason for their high *LDHA* and low *LDHB* expression could be subpar mitochondrial function, especially when compared to other retinal cell types. Thus, we assessed whether there was a mitochondrial activity difference between the photoreceptors and INL cells by examining succinate dehydrogenase (SDH) and cytochrome oxidase (COX) activity in fresh, unfixed, adult retinal sections (Fig. 1l). SDH/complex II plays a role in the citric acid cycle, as well as in the electron transport chain, and its subunits are encoded by the nucleus. COX or complex IV plays a role in the electron transport chain and has catalytic subunits that are encoded by the mitochondrial genome (mtDNA). SDH activity was not lower in the photoreceptors relative to INL cells. COX activity was high in the photoreceptor layer, even higher than that seen in the other retinal layers. The specificity controls for the histochemical reaction are presented in Extended Data Fig. 2i. Finally, IHC for SDH was carried out. The highest IHC signal was observed in the photoreceptor inner segments (IS), as well as the OPL and IPL synaptic layers (Fig. 1m), in good agreement with the observed SDH activity. IHC for another mitochondria-specific enzyme, pyruvate dehydrogenase, showed a similar pattern (Extended Data Fig. 2j) indicating that these are the sites of maximal mitochondrial densities in the retina. Thus, lactate production by the photoreceptors cannot be attributed to lack of mitochondrial activity.

Ldha supports glycolysis by providing a ready supply of cytosolic NAD^+^ that is independent of O_2_ availability and/or mitochondrial function. The phenotype observed following *LDHA* knockdown might be indicative of a reliance on glycolysis where cells might exhibit a preference for unabated and rapid flux through glycolysis. Alternatively, it could be due to an unidentified role of *LDHA* in OS maintenance. To understand the extent of photoreceptors’ dependence on glycolysis, we designed an experimental strategy that satisfied the following criteria: (1) Does not ablate core glycolytic enzymes in order to avoid pleiotropic effects due to their possible non-glycolytic roles, (2) Targets a glycolytic node such that impact on other biosynthetic pathways, such as Pentose Phosphate Pathway (PPP), would be minimal and (3) Uncovers glycolytic reliance and differentiates it from “housekeeping” glycolysis. Glucose-derived metabolites are committed towards glycolytic flux by the enzyme 6-phosphofructo-1-kinase (PFK1), which catalyzes conversion of fructose-6-phosphate (F6P) to fructose-1,6-bisphosphate (F-1,6-BP) (Extended Data Fig. 3a). The most potent allosteric activator of PFK1 is fructose-2,6-bisphosphate (F-2,6-BP)^25^. F-2,6-BP is synthesized from F6P by the kinase activity of the bifunctional enzyme, 6-phosphofructo-2-kinase/fructose-2,6-bisphosphatase (PFK2) (Extended Data Fig. 3a and 3b). To examine the glycolytic dependence of photoreceptors, we targeted the steady state levels of the metabolite, F-2,6-BP as it would satisfy the above criteria.

**Figure 3.**
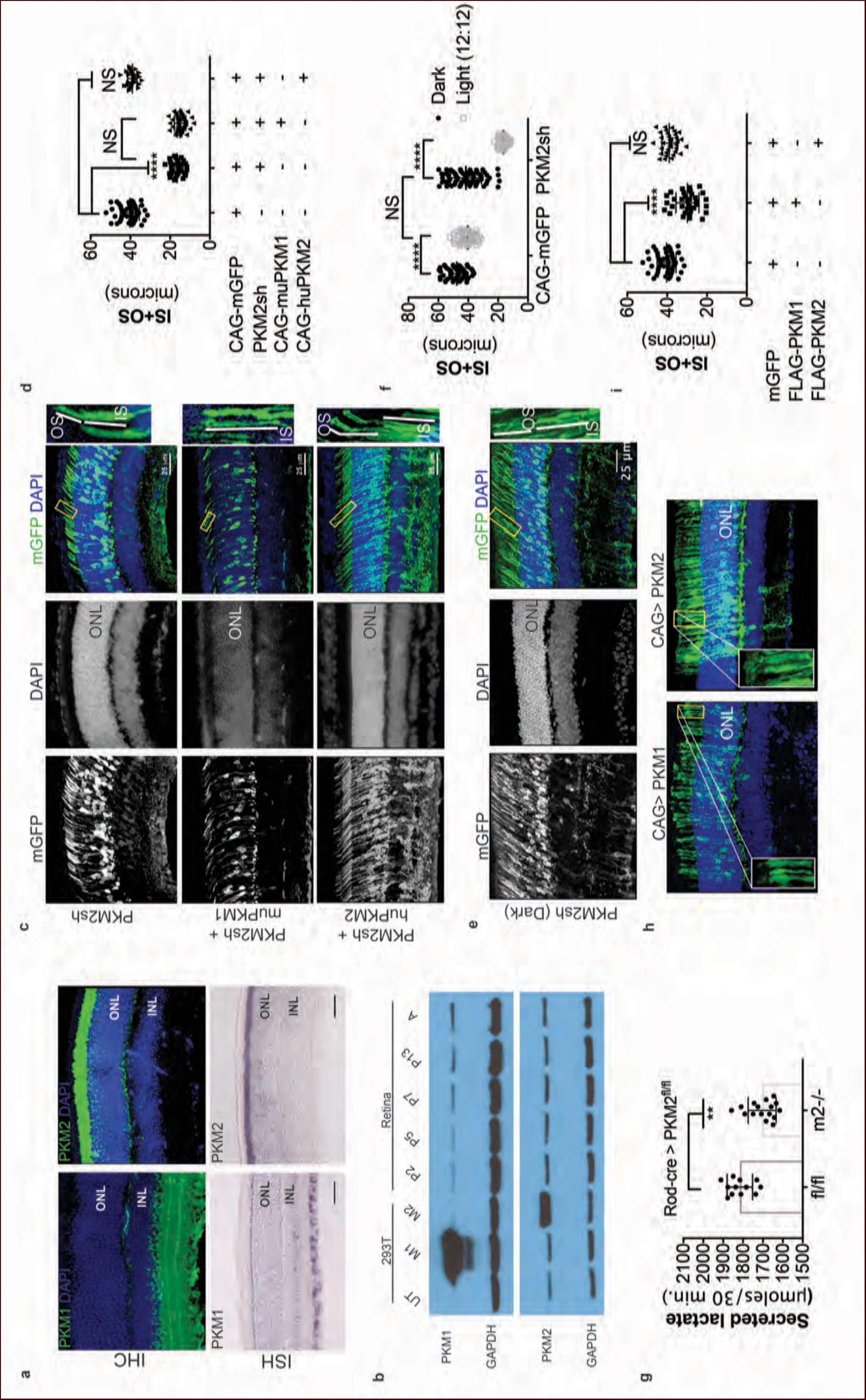
PKM1 and PKM2 isoforms have nonequivalent roles. **a**, Biased expression of M1 and M2 isoforms in retinal layers detected by IHC and ISH. **b**, Immunoblot of retinal lysates from postnatal retina at different developmental stages. HEK293T cell lysates that were from untransfected (UT) cells, or those transfected with CAG-FLAGmuPKM1 (M1) or CAG-FLAGmuPKM2 (M2) as controls. Postnatal age in days. A, mature retina (P25-P30). **c**, Outer segment phenotype of P45 mice after electroporation with constructs encoding mouse PKM2-specific shRNA (PKM2sh) and adding either mouse PKM1 (muPKM1) or human PKM2 (huPKM2). Selected areas in yellow boxes are magnified on the right. **d**, Quantification of IS+OS lengths obtained in (c). n=32-53 cells from 3-4 retinae **e**, Outer segment phenotype of dark-reared P31 mice electroporated with PKM2sh-encoding plasmid. The yellow-boxed region is magnified and presented on the right. **f**, Quantification of IS+OS lengths obtained in (e). n=75 cells from 3 retinae. **g**, Secreted lactate from freshly isolated retinae from PKM2^fl/fl^ (fl/fl) (n= 12) or Rod-cre> PKM2^fl/fl^ (m2^-/-^) (n= 16). **h**, Outer segment phenotype after CAG promoter-driven overexpression of Flag-tagged mouse PKM1 or PKM2. Inset, higher magnification of IS and OS. **i**, Quantification of IS+OS lengths obtained in (h). n=35 cells from 3 retinae in PKM1 and PKM2 groups. ONL, outer nuclear layer. Data, Mean±SD. Statistics, one-way ANOVA with Tukey’s correction for panels **d**, **i**; two-way ANOVA with Tukey’s multiple comparison test for panel **f**; unpaired, two-tailed *t*-test with Kolmogorov-Smirnov correction for panel g.

First we examined expression of PFK2 isoenzymes encoded by *PFKFB1-4* genes (Extended Data Fig. 3c). *PFKFB3* expression could not be detected. *PFKFB1*, *2* and *4* were expressed in either a photoreceptor-enriched or photoreceptor-specific pattern, suggesting a propensity of these cell types to regulate glycolysis via a PFK2-dependent mechanism.

With the exception of Pfkfb3, all other PFK2 isoenzymes have kinase and phosphatase domains on the same polypeptide^26^ (Extended Data Fig. 3b). In addition to potential problems posed by functional redundancy, genetically ablating the PFK2 isoforms would not uncover the preference for directionality. In addition, the structure-function relationships of their kinase and phosphatase domains are not known. To overcome these problems, we overexpressed *TIGAR* (*T*P53-*i*nduced *g*lycolysis and *a*poptosis *r*egulator) as it is functionally similar to the phosphatase domain of PFK2 with well-characterized F-2,6-BPase activity^27^ (Extended Data Fig. 3b) and hence reduces the steady state levels F-2,6-BP. Overexpression of *TIGAR* would be predicted to result in reduced glycolysis without negatively affecting PPP^27^.

We utilized an experimental strategy that addressed the following concerns: (1) The effect should be autonomous to photoreceptors, (2) the phenotype should be induced in fully mature photoreceptors, and (3) the phenotype should discernably be due to perturbations specifically of fructose-2,6-bisphosphate. Our experimental scheme utilized a construct that expressed tamoxifen-inducible Cre only in rods (Fig. 2a, b and Extended Data Fig. 3d). Expression of *TIGAR* specifically in adult photoreceptors resulted in a significant reduction of OS length (Fig. 2c, d). This phenotype was specifically attributable to the phosphatase activity because expression of a catalytic dead version of Tigar, Tigar-<u>TM</u> (triple mutant, H11>A, E102>A, H198>A)^27^, did not cause a change in the photoreceptor OS length (Extended Data Fig. 3e, f). To ascertain if the phenotype is specifically attributable to Tigar phosphatase’s effects on F-2,6-BP, we decided to coexpress *PFKFB3*- a PFK2 isoform that has the kinase activity ~700 fold higher than the phosphatase^28^ (Fig. 2a and Extended Data Fig. 3b). Interestingly, overexpression of *PFKFB3* alone did not result in an overt phenotype- the OS length and morphology were indistinguishable from those of the control electroporated retina (Fig. 2c, d). Overexpression of *PFKFB3* was able to rescue the reduction in OS length caused by *TIGAR* expression (Fig. 2c, d). Together, these data suggest that adult photoreceptors are sensitive to perturbations targeting F-2,6-BP.

Next, the effects of *TIGAR* expression on glycolysis were assayed. Though electroporation answers the question of the cell autonomous effect of a perturbation, the total number of affected cells and their percentage in the electroporated area are too minor to determine biochemical contributions. Adeno-associated virus (AAV)-mediated transduction of TIGAR into photoreceptors was thus used, as it transduces a greater percentage of cells than electroporation. An AAV construct that drives expression of *TIGAR* or mGFP from bovine rhodopsin (*RHO*) promoter specifically in rods was constructed (Fig. 2e, f and Extended data Fig. 3g). The AAVs were injected at postnatal day 6 (P6), after the end of cell proliferation, and nearly full retinal infection was seen by indirect ophthalmoscopy at P24-P27. Retinae were harvested at P28 and examined for expression (Extended data Fig. 3h) and lactate secretion (Fig. 2g). Consistent with the idea that Tigar would interfere in allosteric regulation of glycolysis, a significant reduction in retinal lactate was observed in the AAV-TIGAR infected retinae compared to the control AAV-mGFP infected retinae (Fig. 2g).

Given the essential role of *LDHA* in postmitotic photoreceptors and proliferating cancer cells, other aspects of metabolism that have been discovered in cancer cells, such as the expression of pyruvate kinase isoforms, were investigated. Pyruvate kinase catalyzes the final irreversible reaction of glycolysis and distinct isoenzymes are encoded by two genomic loci, *PKM* (muscle) and *PKLR* (liver and red blood cell). *PKLR* transcripts were not detected in the retina, (data not shown), but M1 and M2 splice isoforms of the *PKM* gene were detected (Extended Data Fig. 4a) in line with protein expression data reported earlier^29^. The M2 isoform is known to regulate aerobic glycolysis, promotes lactate production, and is upregulated in many tumors^30,31^. This isoform was previously reported to be expressed in the photoreceptors^29^. We confirmed that there is photoreceptor-enriched expression of PKM2. PKM1, known to be expressed in most differentiated cell types in adults^32^, was expressed in the cells of the INL and ganglion cell layer, as shown by IHC (Fig. 3a), but was not detectable in photoreceptor cells. In this regard, our data differed from the published findings^29^. To address this discrepancy and validate commercially available antibodies, we performed isoform-specific ISH (Fig. 3a) and confirmed the expression pattern that we observed using IHC. We also examined transcript abundance by qPCR in mRNA purified from isolated rod photoreceptor cells (Extended Data Table 1) and found the M1 isoform to be much less abundant than M2 in the photoreceptors.

**Figure 4.**
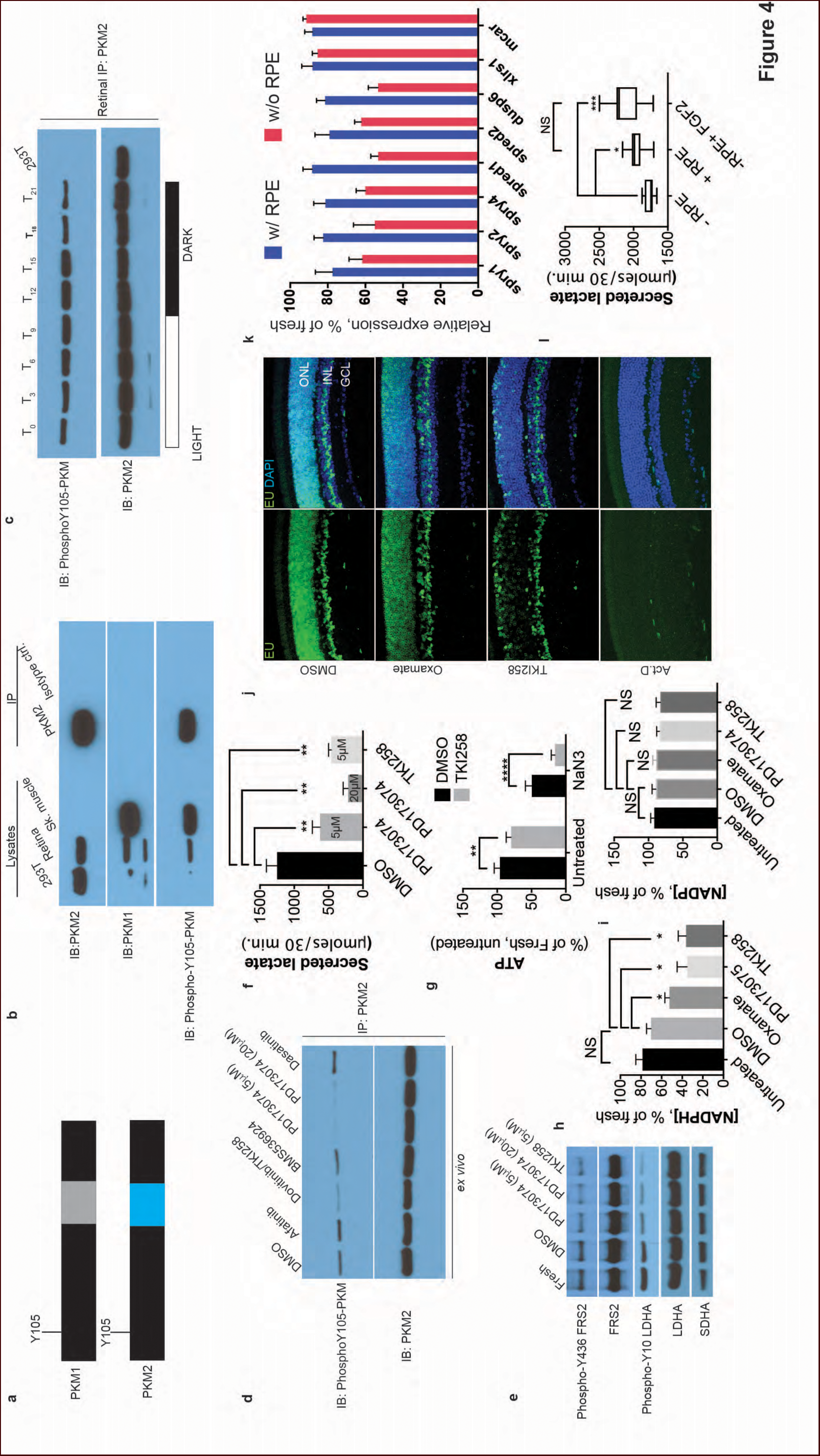
FGF signaling regulates aerobic glycolysis and anabolism **a**, Schematic of PKM1 and PKM2 polypeptide showing Y105 is a shared epitope between PKM1 and PKM2. **b**, Immunoprecipitation (IP) of PKM2 from adult retina followed by immunoblot (IB) for either PKM1, PKM2 or pY105 PKM. IP using isotype-matched antibody is used alongside. Lysates from skeletal muscle (expresses PKM1) and 293T (expresses only PKM2) included as controls. **c**, Retinal lysates were prepared from eyes harvested at 3 hour interval during the 12 hour light 12 hour dark cycle. T_0_ is the time point at light on in the room. The lysates were subjected to immunoprecipitation with anti-PKM2. Immunoprecipitates were probed for phosphorylation at Y105 by immunoblotting with the phospho-specific antibody. **d**, Lysates from explants treated with candidate tyrosine kinase pathway inhibitors or vehicle control (DMSO) were subjected to immunoprecipitation with anti-PKM2. Immunoprecipitates were probed for phosphorylation at Y105 by immunoblotting with the phospho-specific antibody. **e**, FGF inhibitors also reduce phosphorylation of LDHA at the Y10 residue. Phosphorylation of FRS2, an FGFR-interacting protein was included as a control. SDHA served as loading control. **f**, Rate of lactate production from explants treated with DMSO (n=5) or FGF inhibitors PD173074 (5mM) (n=6), PD173074 (20mM) (n=5), TKI258 (n=6). **g**, Steady state ATP levels per retina in explants after culture with TKI258 or DMSO. These retinae were transferred to Krebs’ Ringers with NaN_3_ or NaCl for 30 minutes followed by harvest for ATP extraction. *n*=7, DMSO+NaCl; *n*=9, TKI258+NaCl; *n*=9, DMSO+NaN_3_; *n*=9, TKI258+NaN_3_. Data are Mean±SD. Statistics, Two-way ANOVA with Tukey’s correction. **h**, NADPH steady state levels in explants as a percentage of those measured in freshly isolated retina. Explants were treated with DMSO, oxamate, PD173074, TKI258 or left untreated in culture medium. n=4 groups. Unpaired *t-*test with Kolmogorov-Smirnov correction for indicated pairs. **i**, NADP steady state levels in explants as a percentage of those measured in freshly isolated retina. Explants were treated with DMSO, oxamate, PD173074, TKI258 or left untreated in culture medium. Oxamate, n=5; rest, n=6 groups. Unpaired *t-*test with Kolmogorov-Smirnov correction for indicated pairs. **j**, Blocking glycolysis or FGF signaling reduced EU incorporation in nascent RNA. Explants were treated with DMSO, oxamate, TKI258 or Actinomycin D (RNA Pol II inhibitor) followed by incubation with EU. **k**, Quantitative PCR analysis of transcripts to ascertain relative expression of FGF or non-FGF targets (*MCAR, XLRS1*) in explants cultured with or without RPE/Sclera complex (+RPE or –RPE respectively). **l**, Ability to produce lactate from neural retina increased when cultured in the presence of RPE/Sclera complex (+RPE) (n=11) as compared to those that were cultured without the complex (-RPE) (n=9). Addition of FGF2 in –RPE cultures restored the ability (-RPE+FGF2) (n=8). Retinal explants were cultured with RPE attached in the explant culture medium. Before transferring them to Krebs’s Ringers for lactate estimation, the RPE/Sclera complex was removed and intact neural retina was used. For –RPE conditions, neural retina was cultured in explant medium followed by transfer to Krebs’ Ringers. FGF2 was added to the explant culture medium but was absent in the Krebs’ Ringers for - RPE+FGF2 condition. Data depict median in 1-99 percentile box and whiskers plot. Hinges extend between 25^th^ to 75^th^ percentiles. Statistics, Ordinary one-way ANOVA with Tukey’s correction. ONL, outer nuclear layer. INL, inner nuclear layer. GCL, Ganglion cell layer.

Postnatally, PKM1 expression gradually increased, in correlation with increased differentiation and decreased proliferation in the developing retina (Fig. 3b). On the other hand, PKM2 expression was detectable during the period of proliferation and its expression did not decrease with increased differentiation, likely due to retention of expression in differentiated photoreceptors. Previous studies on pyruvate kinase in the context of proliferation have suggested that loss-of-function of PKM2 reduces proliferation attributable to the glycolytic reliance of mitotic cells for growth^31,33^. To assess if PKM2 plays an essential role in rod photoreceptors, an shRNA construct that specifically targeted mouse PKM2 (PKM2sh), but spared PKM1 (Extended Data Fig. 4b, c, d, e), was generated. *In vivo* electroporation of a plasmid encoding PKM2 shRNA resulted in photoreceptors with significantly shorter OS than control (Figs. 3d, e). This phenotype could be rescued by coelectroporation of a construct encoding human PKM2 cDNA (Fig. 3c, d), which was not targeted by the shRNA (Extended Data Fig. 4f). Coelectroporation of mouse PKM1 with PKM2sh did not rescue the OS length defect (Figs. 3c, d). These data demonstrate that *PKM1* and *PKM2* play nonequivalent roles in rod photoreceptors. We also generated an shRNA construct that targeted exon 4, which is shared between mouse PKM1 and PKM2 (PKM1+2 sh) (Extended Data Fig. 4c, d). Electroporation of this construct resulted in a significant decrease in the OS length (Extended Data Fig. 4g). The photoreceptor morphology and OS length were the same as that observed following electroporation with PKM2sh. While complementation with human PKM2 was sufficient to rescue the IS+OS length defect, we noted some abnormalities with the morphology of some of the photoreceptor IS/OS (Extended Data Fig. 4g). In 4/6 retinae, many photoreceptors lacked clear borders of IS and OS, though in 2/6 retinae, the morphology closely resembled that of control retinae (Extended Data Fig. 4g).

The contribution of PKM2 to OS maintenance was further investigated in the retinae of dark-reared mice electroporated with PKM2sh (Fig. 3e). Dark rearing significantly increased OS length in these animals (Fig. 3f). Taken together, the results from dark-reared animals, in which *LDHA* or PKM2 was knocked down, indicate the requirement for the glycolytic pathway in OS maintenance. Since two different genes that promote aerobic glycolysis are necessary for the light-dependent maintenance of OS, the short OS phenotype is likely due to a reduced supply of the building blocks normally supplied by aerobic glycolysis.

In order to probe the biochemical effects of PKM2 reduction, lactate production was examined. Since electroporated retinae are not ideal for these experiments, mice that had a conditional deletion of PKM2 in rods were used. The PKM2^fl/fl^ mouse strain, in which a M2-specific allele was floxed^33^, was crossed with the Rod-cre strain. The retinae with deficiency of PKM2 had a small but significant decrease in lactate production, as compared to the controls (Fig. 3g). We also noted upregulation of PKM1 in these retinae (Extended Data Fig. 4h) similar to what has been reported before^33^. However, in rods electroporated with PHM2sh, PKM1 expression was not observed (Extended Data Fig. 4i). The presence of the M1-specific exon in the mRNA might reflect a choice made by the splicing machinery after genomic excision of the “preferred” M2-specific exon. This suggestion is made, since a compensatory mechanism would likely have caused PKM1 upregulation after shRNA-mediated knockdown of the PKM2 isoform.

The differential expression of the M1 and M2 isoforms in the retinal layers could be attributable to the differential expression of splicing factors that promote inclusion or exclusion of the M1- or M2-specific exon. To evaluate this possibility, we examined the expression of *SRSF3,* a splicing factor known to promote inclusion of the M2 exon^34^, and *PTBP1,* known to repress the M1 exon inclusion^35^ (Extended Data Fig. 5). While *SRSF3* was expressed at higher levels in photoreceptors, *PTBP1* was more enriched in the INL. Thus the regulation of *PKM* isoform preferences in retina is more complex than that predicted by canonical splicing models.

PKM1 is constitutively active while PKM2 is regulatable^36^. Biased expression of PKM2 in photoreceptors suggests that these cells may need to dynamically regulate glycolysis. The inability of PKM1 to rescue PKM2 loss-of-function indicates that merely replacing pyruvate kinase (PK) function after PKM2 knockdown is not sufficient to restore the OS. In addition, it indicates the importance of glycolytic regulation at the PK step in photoreceptors. We examined the effect of forced expression of PKM1 in the presence of endogenous PKM2, with the hypothesis that the constitutively active isoform might interfere at the regulatory step. We delivered plasmids encoding FLAG-tagged mouse PKM1 and PKM2 via *in vivo* electroporation (Fig. 3h). Photoreceptors electroporated with PKM1-expressing constructs, but not PKM2 expressing constructs, had a reduction in the length of the OS (Figs. 3h, i) with the majority of the photoreceptors in the PKM1 electroporated retinae lacking discernable OS. The two proteins were expressed at equivalent levels, as assessed by western blotting for the FLAG epitope in HEK293T cells (Extended Data Fig. 4j).

PKM2 has been shown to interact with tyrosine phosphorylated proteins^30^ and is tyrosine phosphorylated at position 105 (pY^105^) in tumor cells^37^ leading to promotion of aerobic glycolysis. The pY^105^ is a shared epitope in PKM1 and PKM2 (Fig. 4a). PKM2 was specifically immunoprecipitated from retinal lysates, followed by immunoblotting using a phospho-Y^105^-specific antibody (Fig. 4b). We observed that PKM2 was phosphorylated at Y^105^. In order to ascertain if phosphorylation of PKM2 at this site might have any physiological significance, its regulation by light was examined. PKM2 was immunoprecipitated from the retinae of mice at 3-hour intervals during a 24-hour time course, and phosphorylation at Y^105^ was probed (Fig. 4c). A light-dependent increase in phosphorylation at Y^105^ of PKM2 was observed. Thus, this phosphorylation site might be one of the target sites for physiologically relevant signaling events regulating aerobic glycolysis. This site was then used as a proxy for the tyrosine kinase signaling pathways that could phosphorylate PKM2 in the retina. Freshly explanted retinae were cultured with antagonists targeting specific pathways: Afatinib (EGFR), Dasatinib (Src), BMS536924 (Insulin/IGF), PD173074 (FGFR1) and Dovitinib/TKI258 (FGFR1 and FGFR3). PKM2 was immunoprecipitated and its phosphorylation at Y^105^ probed (Fig. 4d). FGF inhibitors, PD173074 and TKI258, reduced PKM2 phosphorylation. Tyrosine kinase signaling can also target multiple nodes, including pyruvate dehydrogenase kinase and LDHA^38^, and regulate aerobic glycolysis in cancer. We observed that treatment with either PD173074 or TK1258 also resulted in a dose-dependent decrease in Ldha phosphorylation at the Y^10^ residue (Fig. 4e). Thus, FGF signaling potentially targets multiple nodes in order to regulate aerobic glycolysis in the retina.

To determine if FGF signaling might regulate lactate production, freshly explanted retinae were cultured with TKI258 or PD173074, and lactate secretion was measured. Significantly reduced lactate secretion (Fig. 4f) was seen to result from inclusion of either drug. In addition, inhibition of the FGF pathway resulted in increased mitochondrial dependence on ATP steady state maintenance (Fig. 4g). Thus one role for FGF signaling in the adult retina is to promote glycolytic reliance. FGF signaling is required for the maintenance of adult photoreceptors in mice and zebrafish^39–41^. Since it is possible that some of the effects are via regulation of aerobic glycolysis, we examined whether aerobic glycolysis promotes anabolism in the retina. Inhibition of aerobic glycolysis by oxamate treatment or by FGF inhibition resulted in significantly lower steady state NADPH levels-a key cofactor in biosynthetic pathways for lipids, antioxidant responses, and the visual cycle (Fig. 4h). We also observed that interference with aerobic glycolysis did not result in an equivalent reduction in NADP^+^ steady state levels (Fig. 4i). The lowering of NADPH level could be attributable to attenuation of the PPP-shunt as a result of decreased glycolytic flux and/or increased usage of NADPH to quench the reactive oxygen species- an unavoidable consequence of increased mitochondrial dependence. We also assessed other effects on cellular anabolism. Impact of glycolytic perturbation on nucleotide availability was directly visualized by examining nascent RNA synthesis using ethynyl uridine (EU) incorporation. Marked reduction in nascent RNA synthesis was evident following inhibition of Ldh or FGF signaling (Fig. 4i).

Among the large family of FGFs, basic FGF (bFGF/FGF2) has been the most studied in the adult retina. In adult mice and primates, FGF2 is localized to a matrix surrounding photoreceptors and/or is found on their OS^42–44^. The RPE might contribute to a high FGF2 concentration near photoreceptors via biosynthesis, and/or create a barrier to its diffusion from a retinal source. We first examined the role of the RPE in FGF-signaling. Adult retinal explants were cultured with the RPE/choroid/sclera complex, and expression of FGF target genes in the neural retina was compared with that of explants cultured without the attached complex. In the absence of this complex, the transcripts of known FGF signaling targets displayed reduced steady state levels (Fig. 4j). To assess if the reduction in FGF targets was part of a general transcription downregulation or specifically due to dampened FGF signaling, we examined expression of *XLRS1* or cone arrestin (*MCAR*), genes expressed at moderate levels in the retina^45^ (Fig. 4j). These genes were not downregulated in the absence of the RPE complex.

The effect of the RPE complex on aerobic glycolysis was analyzed by quantifying lactate production (Fig. 4k). Culturing retina in the presence of the RPE complex resulted in a small, but significant, increase in the ability to produce lactate. Addition of FGF2 in the culture medium was sufficient to increase lactate production from explants cultured without the RPE. Together these data suggest that the RPE/choroid/sclera complex contributes to FGF signaling in the neural retina and that this signaling pathway plays a role in regulating AG.

Several reports suggest that aerobic glycolysis is a feature of some normal proliferating somatic cells^20,46,47^, and not just of cancer cells. Our work expands the cell types where aerobic glycolysis can occur to include a mature cell type, the differentiated photoreceptor cell. A critical aspect of aerobic glycolysis is its ability to be regulated. The data presented here suggest that allostery and FGF signaling are the regulatory mechanisms in the retina. We favor a model where Aerobic glycolysis appears to be relevant to photoreceptors not only for organelle maintenance, but likely also helps photoreceptors meet their multiple metabolic demands (Extended Data Fig. 6).

Aerobic glycolysis in the retina may have implications for blinding disorders. Studies on retinal degenerative disorders indicate that there are metabolic underpinnings to photoreceptor dysfunction, especially those centering around glucose uptake and metabolism^48,49^. Furthermore, reducing metabolic stress prolongs survival and improves the function of photoreceptors^50,51^. In such treated retinae, there is a trend towards upregulation of glycolytic genes^
51^ or metabolites^52^. However, a direct cause-and-effect relationship between cell survival and glycolysis has not been established. Our results highlight the metabolic strategies employed by healthy photoreceptors and provide a rational basis for the identification of candidate factors that would further clarify the role of glycolysis in retinal degeneration.

## Acknowledgements

We thank Ryan Chrenek, Parimal Rana, Lillian Horin and Alexandra McColl-Garfinkel for technical help. We are grateful to Will Israelsen and Matthew Vander Heiden for help with PKM2^fl/fl^ mice, Ying-Hua Wang and David Scadden for LDHA^fl/fl^ mice and Yun Z. Le for Rod-cre mice. We are indebted to Barry A. Winkler for generous input on retinal metabolism, especially at the earlier stages of this work. We thank Ben Stranges (George Church lab) and Quentin Gilly (Norbert Perrimon lab) for access to the microplate readers. This work has been supported by the National Institutes of Health grant R01 EY023291 and the Howard Hughes Medical Institute.

### Contributions

YC and CC: wrote the paper; YC and CC: conceptualized, designed experiments, interpreted data; YC: designed and generated primary reagents and constructs, standardized explant protocols, performed animal and metabolic experiments; YC and TB: maintained mouse lines, performed expression analysis by western blotting, IHC and ISH; YC and ED: standardized immunoprecipitations; DW: purified RNA from isolated photoreceptors, performed indirect ophthalmoscopy.

### Competing financial interests

Authors declare no competing financial interests.

## Methods

### Plasmids, viruses, *in vivo* electroporation and transfection

The synthetic promoter CAG consisting of cytomegalovirus (CMV) enhancer, chicken β-actin and rabbit β-globin gene splice acceptor was used for expression and genetic complementation. The expression pattern from this promoter when delivered by electroporation has been described previously^1^. Co-electroporation of a plasmid encoding myristoylated/membrane green fluorescent protein (mGFP) allowed visualization of cells that received the plasmid and marked the inner and outer segments. Co-electroporation rate of plasmids to the retina is close to 100%^1^. Full-length rat LDHB (rLDHB), human TIGAR, mouse PFKFB3 and human PKM2 cDNA were obtained from Open Biosystems/GE Dharmacon. Subcloning, epitope tagging and site-directed mutagenesis were carried out by routine molecular biology procedures. For short hairpin (sh) design targeting PKM, and LDHA following resources/software were used: The RNAi consortium, CSHL RNAi central, iRNAi, Invitrogen Block-iT^TM^ RNAi designer. Designed sh oligos were subcloned in pLKO.1 TRC backbone to be driven by the U6 promoter (Addgene, #10878) and the sequences used in this manuscript are listed in Extended Data Table 2. Four hairpin constructs were screened for LDHA and 72 were screened for PKM1/PKM2. Those hairpins that targeted specific mouse sequences but did not target human PKM2 were chosen. The murine FLAG-tagged PKM1 and PKM2 cDNAs were obtained from Addgene (#44240 and #42512) and subcloned in pCAG-EN. The pyruvate kinase activity from these ORFs has been already reported^2^.

The plasmids were mixed in equal molar ratios by accounting for their lengths and subjected to Phenol:Chloroform extraction followed by ethanol precipitation and resuspended to a final concentration of 1 mg/mL in Phosphate Buffered Saline. Subretinal *in vivo* injections and electroporation were carried out as described earlier^3^. For knockdown assays or testing expression from plasmids, transfection in HEK293T cells was carried out as using polyethylenimine (PEI). For making the AAV-mGFP and AAV-TIGAR constructs, the CMV promoter in the empty AAV-MCS8 vector (Harvard Medical School DF/HCC DNA Resource Core) was replaced with the bovine rhodopsin promoter^1^. Woodchuck hepatitis virus posttranscriptional response element (WPRE) was added to enhance expression. Capsid type 8 AAVs were produced and titered as described previously^4^. For subretinal injections of AAV, ~3.5 × 10^6^ – 5 × 10^6^ particles (based on genome copies) per eye were used. P6 pups were injected in order to transduce cells after the proliferative phase of retinogenesis so as to minimize any detrimental effects on cell division and dilution of replication-incompetent viruses. The extent of infection was assessed with a Keeler indirect ophthalmoscope using the cobalt blue filter and Volk 78 diopter lens on non-anesthetisized animals. Mice with edge-to-edge infection were tagged and used subsequently for lactate assays and immunoblotting.

### Mice and animal husbandry

Timed pregnant, wildtype CD1 female mice were obtained from Charles River Laboratories and P0-P1 pups thereof were used in electroporations. C57BL/6J and the two-color Cre reporter mouse *Gt(ROSA)26Sor*^*tm4(ActB-tdTomato,-EGFP)Luo*^*/J* (referred to as mT/mG and described previously^5^) were obtained from the Jackson Laboratories. *LDHA*^*fl/fl*^^6^, PKM2^fl/fl7^, Rod-cre^8^ mice have been described before. Rod-Cre; *LDHA*^*fl/fl*^ and Rod-cre; PKM2^fl/fl^ mouse lines were established. For experimentation, these mice were backcrossed with *LDHA*^*fl/fl*
^ or PKM2^fl/fl^ parents and Cre^+^ and Cre^-^ F1 progeny were used to ensure equivalent allelic copies of the *Cre* transgene, minimum genetic difference and ease of age-matching by using the siblings. Animals were housed at room temperature with 12-hour light and 12-hour dark cycle. Tamoxifen injections were carried out as described previously^1^. For dark rearing, electroporated animals were raised their mothers until P11, when the eyes started to open. Following this, they were transferred to animal housing maintained in darkness until weaning age, when they were weaned and group housed in dark until indicated times for harvest. Water and chow were available *ad libitum.* Animal care was following institutional IACUC guidelines.

### Dissections and adult explant cultures

Wild type, pigmented C57BL/6J mice (JAX) were used for explant cultures since presence or absence of RPE was easily discernable. Adult retinae (P23-P28), freshly enucleated eyes were dissected rapidly in Hanks buffered saline solution (HBSS) (Invitrogen). Extraocular tissue was trimmed off and the cornea and iris were carefully removed. Sclera along with the RPE was gently removed. This was done primarily for two reasons: (1) In our assays the presence of Sclera/RPE complex significantly reduced the efficacy of drug treatments and, (2) secreted lactate was not detected from freshly isolated eyecup with intact sclera. Lens was retained to keep the sphericity of the retina for uniform access to the medium. Explant medium consisted of Neurobasal-A (Invitrogen), 0.2% B27 supplement, 0.1% N2 supplement, 0.1% Glutamax and penicillin/streptomycin (all Invitrogen). Retinae were incubated in freshly prepared explant medium constantly supplied with 95% O2 + 5% CO_2_ (Medical Technical Gases) at 37 °C in a roller culture system (B.T.C Engineering, Cambridge, UK) for indicated times. At the end of incubation period, the lens was removed and the retinae were quickly rinsed with prewarmed Krebs’ Ringers medium (98.5 mM NaCl, 4.9 mM KCl, 2.6 mM CaCl_2_, 1.2 mM MgSO_4_, 1.2 mM KH_2_PO_4_, 26 mM NaHCO_3_, 20 mM HEPES, 5 mM Dextrose) saturated with 95% O_2_. Retinae we again incubated in 0.5 mL Krebs’ Ringers medium for 30 minutes in roller culture with 95% O_2_ supplied. The supernatant and retinae were rapidly frozen separately at the end of the experiment. DMSO was used as vehicle control for water insoluble solutes. Sodium oxamate or sodium azide was dissolved in the medium. Equimolar amount of sodium chloride was used as control for osmotic pressure, a colligative property. For +RPE experiments, the extraocular tissue was trimmed off, cornea and iris removed and the eyecups were incubated in the explant culture medium. At the end of the incubation, the RPE/sclera complex was removed along with the lens and the neural retina was incubated in the oxygenated Kebs’s Ringers medium for 30 minutes as described earlier to assay secreted lactate. Thus, our experiments assess the effect of RPE/sclera complex on the ability to produce lactate by neural retina.

### Drugs

Sodium Azide (20 mM, Sigma-Aldrich), Sodium Oxamate (50mM, Sigma-Aldrich), FX11 (10 μm, Calbiochem), BMS 536924 (5 μm, Tocris), Afatinib (5 μm, Selleckchem), Dovitinib/TKI258 (5 μm, Selleckchem), Dasatinib (5 μm, Selleckchem), PD173074 (5 μm or 20 μm, Selleckchem), Actinomycin D (5 μm, Sigma-Aldrich), FGF2 (2 g/mL, Cell Signaling).

### Immunoprecipitation and immunoblotting

BL/6J retinae without RPE were homogenized in Lysis buffer (5 mM HEPES, 1mM DTT, 1mM ATP, 5 mM MgCl2, 1% glycerol, Complete Protease Inhibitor (Roche) and PhosStop phosphatase inhibitor (Roche). Immunoprecipitation was carried out using rabbit anti-PKM2 and rabbit IgG isotype control followed by sheep anti-rabbit-conjugated Dynabeads (Life Technologies). Immunoprecipitates were boiled and loaded on 10% SDS-PAGE gels followed by transfer on Hybond nitrocellulose membranes (GE Amersham). Membranes were blocked with 5% non-fat milk in 1X Tris Buffered Saline + 0.1% Tween-20. A conformation-specific mouse-anti rabbit secondary and HRP-conjugated goat-anti-mouse (Jackson, 1:10,000) tertiary antibodies were used followed by Enhanced Chemiluminescent (ECL) detection using substrate from GE Amersham.

### Immunohistochemistry

Enucleated eyes were fixed overnight at 4°C in 4% formaldehyde. The eyes were passed through an increasing concentration of sucrose (5%, 15%, 30%) followed by equilibration in 1:1 30% sucrose: OCT (Tissue-Tek)and frozen on dry ice. Eighteen micron cryosections were cut using a Leica CM3050S cryostat. Antibodies used are listed in Extended Data Table 3. Heat-mediated antigen retrieval at pH 8 was carried out. For HRP staining, Cell and Tissue staining kit (R&D systems) was used. Confocal images were acquired on Zeiss LSM10 inverted microscope. The intensity and pixel saturation were calibrated for inner and outer segment label (mGFP) so that details in these cellular features were retained. Thus, due to intense signal of mGFP in the outer segments, the labeling in other cells of the inner retina seems variable and less bright despite electroporation known to target these cells. Images were processed on ImageJ. Maximum intensity projections are depicted. Colocalization was confirmed by individual merges of coplanar sections along the z-axis. For IS/OS length measurements, the orthogonal projections of sections were used. The projections spanning the entire IS/OS volume ensure changes due sectioning angle have a minimal effect. Multiple quantifications across the electroporated field were done for at least 3 retinae. Expression by IHC was confirmed in both CD1 (albino) and BL/6J (pigmented) mice. Sclera and RPE were preserved in electroporated eyes to ensure that outer segments were not ripped during the dissections.

### *In situ* hybridization

*In situ* hybridization was carried out as described earlier^9^. Probe sequences are available on request. For *PFKFB1, PFKFB2, PFKFB4, SRSF3* and *PTBP1*, tyramide amplification (Perkin) was used. Bright-field images were acquired on Nikon Eclipse E1000 microscope.

### ATP, Lactate and NADPH assay

For ATP estimation, individual retinae were rapidly frozen in liquid nitrogen at the end of the assay. ATP was measured using ATP bioluminescence kit CLS II (Roche/Sigma-Aldrich). For secreted lactate estimation the retinae were incubated in Krebs’ Ringers medium after indicated treatments. The supernatant from above was used with Lactate assay kit (Eton Bioscience). Amount of lactate produced in 30 minutes was assayed. Intracellular lactate was estimated for AAV-transduced retinae because a large number of mice had to be injected and screened for complete, edge-to-edge infection. Thus, infected retinae at specific age were harvested and frozen as they became available. Two to three retinae were pooled into a group and frozen together. Five such groups (*n*=5) were used for assaying lactate after AAV-mGFP and AAV-TIGAR infection. The retinae were homogenized with the Lactate Assay buffer (Fluorometric Lactate Assay kit, abcam). A small aliquot was removed for protein estimation and subsequent immunoblotting and the remainder was passed through 10 kDa protein filtration column (abcam) to remove proteins and thus minimize interference due to endogenous lactate dehydrogenase in the lactate assay. For protein estimation, Qubit protein assay (Invitrogen) was used since it is not affected by the presence of detergents in the Lactate Assay buffer. NADP and NADPH was assayed using Fluoro NADP/NADPH kit (Cell Technology) following manufacturer’s instructions. The quantifications for NADP and NADPH were made separately and thus represent different retinae and treatments.

### COX and SDH histochemistry

Histochemistry on fresh and unfixed retinal tissue was carried as described earlier for brain tissue^10^. The assay relies on the ability of functional cytochrome oxidase to catalyze oxidative polymerization of 3,3’-diaminobenzidine (DAB) (an electron donor) to brown indamine product. Succinate dehydrogenase assay is based on the ability of this enzyme to oxidize supplied succinate and in turn reduce a ditetrazole (NBT) to dark blue diformazan using phenazine methosulfate (PMS), an intermediate electron carrier.

### 5-ethynyl uridine (EU) labeling

Explants were cultured with indicated drug or DMSO for 5 hours followed by 1mM EU (Life Technologies) with the drug for additional 2.5 hours. The retinae were fixed, cryosectioned and processed for label detection using Click chemistry reagents (Life Technologies).

### Quantitative RTPCR

RNA was isolated using TRIzol reagent (Life Technologies) from 3-4 retinae. Two μg RNA was subjected to cDNA synthesis using SuperScript III reverse transcriptase and random hexamers. QPCR was performed using power SYBR Green PCR Master mix (Applied Biosystems) on a 7500 Fast Real-Time PCR System (Applied Biosystems). Primer sequences are provided in Extended Data Table 4. *RPL13a* was used as internal reference and freshly isolated retinal tissue was used as calibrator sample. Expression ratio was calculated using 2^-ΔΔCt^ method. For each target gene, three technical replicates were simultaneously assayed to arrive at the average value for a biological replicate. Mean of three biological replicates was used to derive the C_t_ value of each target.

### Rod isolation and cDNA synthesis

P0 CD1 mice were electroporated with Rho-dsRed plasmid which encodes for dsRed, driven by bovine rhodopsin promoter, which results in retinas with patches of dsRed expression only in rod photoreceptors^11^. Once they reached adulthood, mice were then euthanasized via CO_2_ asphyxiation and the retinas were rapidly removed. The retinas were incubated for 5 minutes at 37**°** C in Hank’s Balanced Salt Solution (HBSS) supplemented with 10 mM HEPES and 5 mM EDTA and then gently triturated with a P1000. The dissociated retina was allowed to settle on sylgard-coated petri dishes. Rods expressing the dsRed reporter were identified by their red fluorescence using an inverted microscope and hand-pipeted directly into lysis buffer, and their cDNA amplified using the previously described protocol that utilizes oligo dT priming^12^.

### Data collection and statistics

Data collection was from non-randomized experiments. The primary experimenters were not blinded to treatments. No statistical methods to predetermine sample size were employed. No assumptions for potential outliers were made and hence all data points were included in analyses and depicted. Normality of data distribution was tested using D’Agostino-Pearson omnibus test. Non-parametric statistics were used when Gaussian distribution of data points could not be obtained. *p-*value denoted as: Not significant (NS), *p* > 0.05; *, *p* ≤ 0.05; **, *p* ≤0.01; ***, *p* ≤0.001; ****, *p* ≤0.0001.

